# Formate-induced CO tolerance and innovative methanogenesis inhibition in co-fermentation of syngas and plant biomass for carboxylate production

**DOI:** 10.1101/2022.06.30.498223

**Authors:** Flávio C. F. Baleeiro, Lukas Varchmin, Sabine Kleinsteuber, Heike Sträuber, Anke Neumann

**Affiliations:** Department of Environmental Microbiology, Helmholtz Centre for Environmental Research – UFZ, Leipzig, Germany; Technical Biology, Institute of Process Engineering in Life Science, Karlsruhe Institute of Technology – KIT, Karlsruhe, Germany

**Keywords:** Mixotrophy, Volatile fatty acids, Medium-chain carboxylic acids, Formic acid, Chain elongation, Microbiome, Carbon monoxide, Ethene, Hexanoic acid, Lactic acid bacteria

## Abstract

Production of monocarboxylates using microbial communities is highly dependent on local and degradable biomass feedstocks. Syngas or different mixtures of H_2_, CO, and CO_2_ can be co-fed to a fermenter to alleviate this dependence. To understand the effects of adding these gases during anaerobic fermentation of plant biomass, a series of batch experiments was carried out with different syngas compositions and corn silage (pH 6.0, 32°C). Co-fermentation of syngas with corn silage increased the overall carboxylate yield per gram of volatile solids (VS) by up to 44% (0.36 ± 0.07 g g_VS_^-1^; in comparison to 0.23 ± 0.04 g g_VS_^-1^ with a N_2_/CO_2_ headspace), despite slowing down biomass degradation. Ethylene and CO exerted a synergistic effect in preventing methanogenesis, leading to net carbon fixation. Less than 12% of the electrons were misrouted to CH_4_ when either 15 kPa CO or 5 kPa CO + 1.5 kPa ethylene was used. CO increased the selectivity to acetate and propionate, which accounted for 86% (electron equivalents) of all products at 49 kPa CO, by favoring lactic acid bacteria and actinobacteria over *n*-butyrate and *n*-caproate producers. This happened even when an inoculum pre-acclimatized to syngas and lactate was used. Intriguingly, the effect of CO on *n*-butyrate and *n*-caproate production was reversed when formate was present in the broth. The concept of co-fermenting syngas and plant biomass shows promise in two aspects: by making anaerobic fermentation a carbon-fixing process and by increasing the production of propionate and acetate. Testing the concept in a continuous process could improve selectivity to *n*-butyrate and *n*-caproate by enriching chain-elongating bacteria adapted to CO and complex biomass.

## BACKGROUND

Anaerobic fermentation enables the production of carboxylates from cheap organic matter. The technology uses microbial communities as robust and self-regenerating biocatalysts to produce a spectrum of carboxylates with one to eight carbon atoms in their chain [1, 2]. Anaerobic fermentation is basically a methane-arrested anaerobic digestion [1], hence, it can take profit from the existing infrastructure and knowledge about anaerobic digestion while producing chemicals that are more valuable and energy-dense than biogas [3]. Carboxylates, in particular those with longer carbon chains, can be extracted from fermentation broths with relative ease [4] and hold promise as a bio-based platform for the chemical and fuel industry [5].

Ultimately, the impact that anaerobic fermentation can have as a sustainable technology for transforming the economy is limited by the availability of organic feedstocks that are local, cheap, and biodegradable [6, 7]. Regardless if supplied by industrial waste streams, municipal organic waste, or plant biomass, the feasibility of the anaerobic fermentation plant is constrained by biomass cost, supply, and quality.

One way to make anaerobic fermentation more independent of organic feedstocks is by exploiting mixotrophic microorganisms that are highly efficient in consuming both organic and inorganic substrates [8]. Thanks to the reductive acetyl-CoA pathway present in many anaerobic bacteria, mixtures of H_2_, CO_2_, and CO (often called syngas) are one of the most promising inorganic co-substrates for anaerobic fermentation [9, 10]. These gases allow to merge anaerobic fermentation of biomass with other promising green technologies such as dry biomass gasification (via syngas), power-to-gas (via H_2_), carbon capture (via CO_2_), or treatment of industrial off-gases (via CO) [11]. There are numerous ways to combine such technologies [12], and fermenting syngas and complex biomass in a one-pot process is one of the simplest [5].

Recently, several studies have explored the use of microbial communities to produce carboxylates from syngas components in a one-pot process. They approached the topic by 1) innovative ways of supplying H_2_ [13, 14], 2) enrichment of microbial communities in mineral media with syngas [15-17], 3) co-fermentation of H_2_/CO_2_ (or H_2_/CO_2_/CO) with synthetic organic substrates [18-20], and 4) co-fermentation of H_2_/CO_2_ with organic waste streams [21]. In general, the presence of H_2_ and CO helped increasing carboxylate yields and selectivity to medium-chain carboxylates. Yet, we found no studies with complete syngas mixtures (H_2_/CO_2_/CO) and plant biomass.

CO is expected to have a major impact on the plant biomass fermentation. Despite being a key substrate for acetogenic bacteria, carbon monoxide is a strong inhibitor of hydrogenases [22] and hence disrupts the electron transport chains of bacteria that rely on these enzymes. This is aggravated in fermentative bacteria, which often rely on [Fe-Fe] hydrogenases that are particularly sensitive to CO [23]. Yet, there are ways to overcome CO inhibition of fermentative bacteria. For instance, *Clostridium kluyveri* was less prone to CO inhibition when grown in co-culture with acetogens (e.g., *Clostridium autoethanogenum*) in bottles without shaking [24]. Besides, the presence of formate, a common extracellular metabolite in anaerobic microbial communities, allowed *Acetobacterium woodii* to tolerate up to 25 kPa CO [25], and low concentrations of formate (<100 mg L^-1^) improved the growth of *Clostridium ljungdahlii* and *Clostridium carboxidivorans* on syngas [26].

In this study, we aimed to understand the main effects of syngas on the carboxylate production during the anaerobic fermentation of plant biomass by using corn silage as a model feedstock. We focused on the syngas composition taking in consideration the double-edged role of CO in carboxylate production.

## MATERIAL AND METHODS

### BATCH CULTIVATION

In an anaerobic chamber, 50 mL of anoxic mineral medium (preparation and composition are described in Additional File 1 and Table S1, respectively) were added to 250-mL-serum bottles containing 5.5 g of fresh matter of corn silage each, resulting in a VS (volatile solids) concentration of 30 gVS L-1. All bottles contained at least the autochthonous microbial community of the corn silage as inoculum. Bottles additionally inoculated with a community adapted to syngas received cells from an enrichment reactor (see “Source of the adapted community”). To harvest the cells from the enrichment reactor, the fermentation broth was centrifuged at 4,816 × g and 4°C for 10 minutes. The supernatant was discarded and the cells were washed and resuspended in the equal volume of anoxic mineral medium. Alternatively, the untreated broth from the enrichment reactor, which was rich in carboxylates, was either used instead of mineral medium or 10% of the medium was substituted with it (inoculation with 100% or 10% reactor broth). The serum bottles were closed with butyl rubber stoppers and sealed with aluminum crimps. All batch cultures were prepared in duplicates. The bottles were initially pressurized with 150 kPa (1.5 bar_a_). Different partial pressures of H_2_ and CO were applied (Fig. 1), resulting in a combined partial pressure of 98 kPa. This ensured a fixed amount of electron donors in the headspace. The only exceptions were the condition “CO_2_”, which contained neither CO nor H_2_, and the condition “CO+CO_2_^”^, which contained 49 kPa CO but no H_2_. All cultivation bottles initially contained 24 kPa CO_2_. N_2_ was used as a filling gas to reach the final pressure. In two pairs of bottles, 3 mL ethylene was additionally added to the pressurized bottles to achieve 1.5 kPa ethylene at 0 kPa CO (98 kPa H_2_) and 5 kPa CO (93 kPa H_2_) (Fig. 1). In one pair of bottles, 0.2 mL of formic acid was added at 9 kPa CO (89 kPa H_2_) to reach a concentration of 5 g L^-1^ formate (Fig. 1). The pH was corrected with 4 M KOH immediately after formic acid addition. For abiotic controls, the sealed and pressurized bottles with corn silage and mineral medium were autoclaved for 20 minutes at 121°C.

**Fig. 1.**
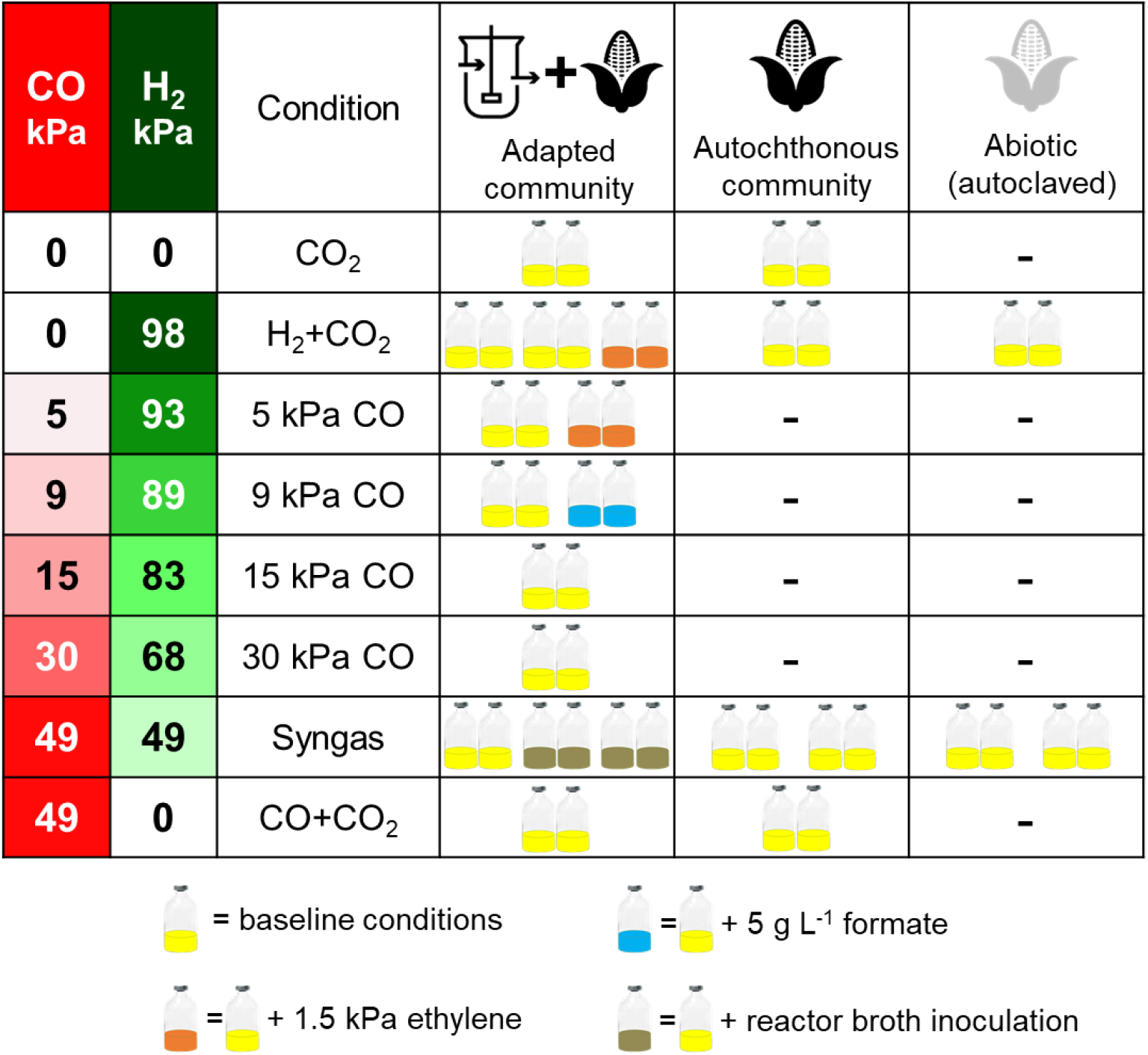
Summary of all conditions tested in batch cultures. Yellow bottles contained at least corn silage, mineral medium, and CO_2_. Additionally, CO and H_2_ were added as indicated. As special conditions, formate or ethylene was additionally added, or untreated reactor broth was used as source of the adapted community instead of washed cells.

The batch cultures were incubated on a rotary shaker at 32°C and 200 rpm. Fermentations were carried out at a pH value of 5.9 ± 0.5. Sampling of the liquid phase and pH correction with 4 M NaOH were done weekly. The pressure was monitored twice per week. When a pressure lower than 100 kPa was detected, the bottle was re-pressurized to 150 kPa. Before every pressurization, the headspace of the bottle was purged with 2.5 L of the corresponding gas mixture. By monitoring the chemicals in the gas and aqueous phase, little to no activity was detected in the abiotic controls (Figure S1), and the amount of DNA extracted from cell pellets was too low for microbial community analysis. For practical reasons, the experiments were divided in three different batches lasting from 31 to 38 d. For information on how the experiments were divided in different batches and the exact duration of each batch see “Further details on the experimental setup” in Additional file 1.

CORN SILAGE. Corn silage from a farm in Neichen (Saxony, Germany) was transported and stored in vacuum-sealed polyester bags at room temperature until its use. Soluble chemicals in the silage were quantified (0.41 ± 0.04 g L^-1^ acetate, 0.21 ± 0.06 g L^-1^ ethanol, and 3.2 ± 0.1 g L^-1^ lactate). The content of VS on a fresh matter basis for the substrate was 27.1 ± 0.1%.

### SOURCE OF THE ADAPTED COMMUNITY

The syngas-adapted community used as inoculum for the batch cultures originated from two 1.0-L stirred-tank reactors that were being operated near atmospheric pressure (102 kPa) with continuous syngas recirculation (32% H_2_, 32% CO, 16% CO_2_, 2% ethylene, 2% He, and rest N_2_ at 40 mL min^-1^) at a hydraulic retention time of 14 d, pH 6.0, and 32°C. Once a day, fermentation broth was harvested and mineral medium (Table S1) containing additionally 12 g L^-1^ acetate and 12 g L^-1^ lactate was fed. Other reactor operation procedures were done as described by Baleeiro, et al. [20]. At the time the inoculum was collected, the enrichment reactors had been operated for 83 days and had an average carboxylate composition of 11 g L^-1^ acetate, 3.4 g L^-1^ *n*-butyrate, 2.6 g L^-1^ *n*-caproate, and 1.1 g L^-1^ *i*-butyrate. Methanogenesis rates were very low (about 2.2 mL CH_4_ L^-1^ d^-1^) and syngas consumption rates were relatively high (150 mL H_2_ L^-1^ d^-1^ and 237 mL CO L^-1^ d^-1^ on average). The original microbial community composition of the adapted community is shown in Figure S2.

### ANALYTICAL METHODS

The VS content of corn silage was determined according to Strach [27]. To estimate the corn silage degradation during fermentation, a procedure based on total solids (TS) measurement was applied. At the end of the fermentation, the whole content of each bottle was separated by sieving (0.84 mm mesh size). The retained solids were washed with 50 mL phosphate buffer saline solution (PBS; pH 7.4, 11.8 mM phosphates) for 30 minutes at room temperature in a rotary shaker at 150 rpm. Afterwards, the resulting mixture was sieved again and the TS content of the solid fraction was determined by drying it at 105°C until a constant weight was obtained (24 to 48 h). The difference between the TS content (in %) of the abiotic control and the TS content (in %) of the test bottle was defined as the solids degradation, in percentage points (p.p.).

For measuring pH and concentrations of chemicals in the bottles, 0.3 mL of liquid was collected once a week from each bottle. Liquid samples were centrifuged at 9,000 × g for 10 minutes and a defined amount of the resulting supernatant was diluted five times with PBS to reach a neutral pH value before being analyzed. For analyzing the concentration of chemicals in corn silage, an elution of 25 g substrate with 250 mL PBS was carried out for 24 h at room temperature [28]. The resulting liquid samples were filtered with 0.22 µL nylon syringe filters before being analyzed via high performance liquid chromatography (HPLC). Concentrations of linear monocarboxylates from formate (C1) to *n*-caprylate (C8), linear alcohols from ethanol (C2) to *n*-hexanol (C6) as well as lactate, *i-*butyrate, *i*-valerate, and *i*-caproate were measured using a 1100 series HPLC-RID/UV system (Agilent Technologies, Germany) equipped with pre-column and column Rezex ROA-Organic Acid H+ (8%) (Phenomenex, Germany). For compounds with UV absorption at 280 nm, concentration values obtained via UV were averaged with the values obtained via a refractive index detector (RID). The mobile phase of HPLC was 5 mM H_2_SO_4_ at 0.6 mL min^-1^. A sample injection volume of 20 µL was used and the temperatures of the column oven and RID were kept at 55°C and 50°C, respectively. Concentrations of *n-*propanol, *n-*butanol, *n-*pentanol, *n-*hexanol, *n*-heptanoate, *n*-caprylate, *i*-valerate, and *i*-caproate in this study generally remained below 100 mg L^-1^ and are not shown.

The bottle headspace pressure was analyzed using a manometer GDH 14 AN (Greisinger electronic, Germany). Afterwards, 2 mL of gas sample was collected twice a week for gas composition analysis via gas chromatography (GC) with a thermal conductivity detector (TCD). The GC-TCD system measured the fractions of H_2_, CO_2_, CO, ethylene, N_2_, O_2_, and CH_4_ and was described previously by Mohr, et al. [29].

Rates in this study are average rates specific to the volume of broth (50 mL). Average rates in terms of electron equivalents were used to compare production and consumption of chemicals under different cultivation conditions. Conversion factors in Table S2 were used for converting mass values into moles, electron equivalents, and carbon equivalents when necessary.

### MICROBIAL COMMUNITY ANALYSIS

For microbial community analysis, 16S rRNA amplicon sequencing was performed using the Illumina MiSeq platform. Cell pellets were collected from the culture bottles at the end of the experiments. Before being stored at -20°C, cell pellets were washed once with an equal volume of PBS (ca. 1.8 mL). DNA was extracted from the cell pellets using the ZymoBIOMICS DNA Miniprep kit (Zymo Research, Germany) with cell disruption within 20 min, following the manufacturer’s instructions for non-soil samples. DNA quantification and quality assessment, polymerase chain reaction (PCR), and library preparation were done as described by Logroño, et al. [30] for 16S rRNA. For PCR, primers for the V3 and V4 regions [31] were used. Filtering, denoising, and taxonomical assignment of the amplicon data were done as described previously [32]. Sequence counts of all samples were rarefied to the read number of the sample with the lowest coverage in the dataset (56,674 counts).

Raw sequence data for this study was deposited at the European Nucleotide Archive (ENA) under the study accession PRJEB49567 (http://www.ebi.ac.uk/ena/data/view/PRJEB49567).

## RESULTS AND DISCUSSION

### COMPARING DIFFERENT STARTING CONDITIONS

To assess the impact of using a microbial community previously adapted to syngas and lactate (besides the autochthonous corn silage community), we compared inoculated cultures with cultures containing only the corn silage community. Figure S3 presents the product profiles of the fermentations with syngas (49 kPa CO, 49 kPa H_2_, 24 kPa CO_2_) depending on the way the adapted community was inoculated. As long as the inoculum did not contain high amounts of carboxylates from the enrichment reactor, i.e. when the inoculation was done with washed cells or with 10 vol % reactor broth, using the adapted community proved to be particularly important for accelerating the consumption of H_2_ and CO. Yet, the maximum *n-*butyrate concentration was 1.2 ± 0.1 g L^-1^ and the adapted community contributed little to produce carboxylates longer than propionate. Since the addition of reactor broth at the beginning of the fermentations was not beneficial to produce medium-chain carboxylates, only washed cells were used in further tests with the adapted community.

The results of this preliminary experiment (Figure S3) are summarized in the Additional file 1 (“Inoculation with an adapted community”).

### EFFECTS OF CO, H_2_, AND SYNGAS

Inoculating a community adapted to syngas was not enough to allow medium-chain carboxylate production from syngas and corn silage. Therefore, we tested the effects of the main syngas components (H_2_ and CO) separately on the fermentation with both the autochthonous and the adapted communities.

Fig. 2 shows the production and consumption rates of chemicals (in electron equivalents) together with the community composition under each condition. In cultures with CO, regardless of the community type, propionate was a main electron sink, whereas cultures without CO mostly routed more electrons to *n*-butyrate (Fig. 2a). The presence of CO (49 kPa) inhibited the production of carboxylates with chains longer than propionate by both communities.

**Fig. 2.**
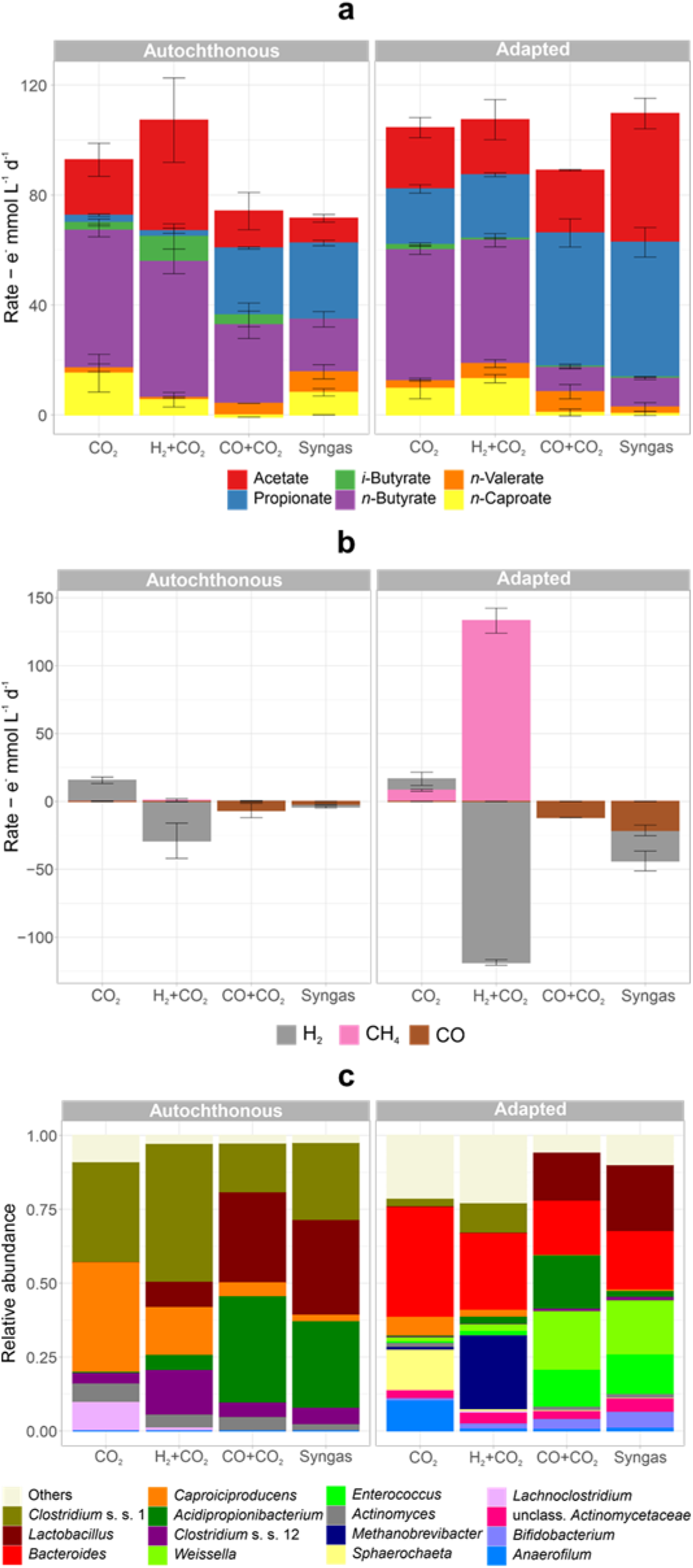
Production and consumption rates for liquid (**a**) and gaseous (**b**) chemicals and microbial community composition (**c**) in fermentations of corn silage with different syngas constituents. The 15 most abundant genera in the set are shown. Mean values of duplicate bottles are shown. Error bars indicate standard errors. Rates are in terms of electron equivalents. S.s.: sensu stricto.

Methane formation occurred only in the culture with the adapted community and was stronger when H_2_ was present and CO was absent (H_2_+CO_2_, Fig. 2b). In this case, routing of electrons to CH_4_ occurred to a similar extent as H_2_ consumption (133 ± 9 e^-^ mmol CH_4_ L^-1^ d^-1^ and 119 ± 2 e^-^ mmol H_2_ L^-1^ d^-1^), indicating the activity of hydrogenotrophic methanogens in the adapted community. This was confirmed by the high relative abundance of *Methanobrevibacter* under this condition (H_2_+CO_2_ with the adapted community, Fig. 2c).

The autochthonous community alone consumed some exogenous H_2_ and produced extra acetate when CO was absent (H_2_+CO_2_, Fig. 2b). Still, the adapted community achieved the highest CO consumption (21 ± 4 e^-^ mmol CO L^-1^ d^-1^), which occurred when both H_2_ and CO were present (Syngas, Fig. 2b). Under this condition, the adapted community presented a net carbon fixation rate of 3.8 ± 0.7 C mmol L^-1^ d^-1^ (equivalent to 167 ± 31 mg CO_2_ L^-1^ d^-1^) (Figure S4).

H_2_ and CO were likely converted to acetate by autotrophic acetogens such as *Clostridium* sensu stricto 12, a genus that was present (Fig. 2c) and comprises acetogenic species. However, cultures with the adapted community with the highest H_2_ and CO consumptions (excluding methanogenesis) had low relative abundances of *Clostridium* sensu stricto 12. The highest relative abundances were recorded for bacteria likely growing heterotrophically on sugars and lactate from the plant biomass: lactic acid bacteria (LAB; *Lactobacillus, Weissella, Enterococcus*, and *Bifidobacterium*), *Bacteroides, Acidipropionibacterium, Actinomyces*, and some clostridia (*Clostridium* sensu stricto 1, *Caproiciproducens*, and *Anaerofilum*).

The presence of H_2_ or CO disadvantaged *Lachnoclostridium* (autochthonous community) and *Sphaerochaeta* and *Anaerofilum* (adapted community) (Fig. 2c), genera commonly associated with improved lignocellulose degradation [33-36]. Other major shifts in microbial composition were due to the presence of CO. Cultures with CO (49 kPa CO) had greater relative abundances of LAB and *Acidipropionibacterium* at the cost of *Caproiciproducens, Methanobrevibacter*, and *Clostridium* sensu stricto 1. Swaps of relative abundances of lactate- and *n*-butyrate-producing bacteria also occur in other anaerobic systems and depend on pH and organic substrate type [37]. Cultures without inoculation with syngas-adapted cells showed higher relative abundances of *Caproiciproducens, Clostridium* sensu stricto 1 and 12, *Actinomyces*, and *Acidipropionibacterium*, whereas cultures inoculated with the adapted community tended to have higher shares of *Bacteroides, Enterococcus, Weissella*, and *Bifidobacterium. Weissella, Bifidobacterium*, and *Enterococcus* were not detected in the adapted community inoculum, indicating that they originated from the corn silage community. *Bacteroides* was present in low abundances in the adapted community inoculum (Figure S2).

Propionate production by the autochthonous community in the fermentations with CO (Fig. 2a) can be explained by high abundances of *Acidipropionibacterium* (Fig. 2c), which compete with clostridia for lactate [38]. Yet, the adapted community produced even more propionate under the same conditions (Fig. 2a), despite low abundances of *Acidipropionibacterium*. So far, no isolated carboxydotroph is known to convert CO to propionate, but co-cultures of carboxydotrophs and propionate producers can [39]. Thus, indirect conversion of CO to propionate intermediated by ethanol is possible. However, high abundances of heterofermentative LAB (i.e. *Weissella* and *Lactobacillus*, which can produce propionate from sugars and lactate [40, 41]) strongly indicates that electrons in propionate originated from the plant biomass and not from CO.

Interestingly, the relatively high H_2_ and CO consumption by the adapted community in the fermentations with syngas (45 e^-^ mmol L^-1^ d^-1^ of gases consumed and 109 e^-^ mmol L^-1^ d^-1^ of carboxylates produced) did not result in much higher carboxylate production than that when only CO_2_ was in the headspace (carboxylate production of 105 e^-^ mmol L^-1^ d^-1^) (Fig. 2a). This observation suggests that less plant biomass was consumed when H_2_ or CO were present, which was confirmed by the analysis of the degradation degree (Figure S5). The presence of H_2_ alone also slowed down biomass degradation, especially by the communities without methanogenic activity (autochthonous community, Figure S5). Headspaces rich in H_2_ and H_2_+CO_2_ have been reported to slow down the degradation of organic waste streams. Arslan, et al. [42] noted a lower hydrolysis rate under a H_2_-rich headspace, but not a change in the final hydrolysis degree of the solid fraction, implying a retarded utilization of the available organic feedstock. There, H_2_ and H_2_+CO_2_ increased the overall carboxylate production by 47% and 150%, respectively, in comparison to a fermentation with a N_2_ headspace.

Within the same community type (i.e. adapted or autochthonous), community compositions of cultures fed with CO (conditions “Syngas” and “CO+CO_2_”) were very similar. This fact pointed to the importance of CO among all other factors. To get a more detailed view on the effect of CO on the fermentation, further tests with partial pressures lower than 49 kPa CO were carried out.

### EFFECT OF DIFFERENT CARBON MONOXIDE CONCENTRATIONS

Production rates for chemicals and community compositions at partial pressures between 0 and 49 kPa CO for cultures with the adapted community are shown in Fig. 3. During this experiment, partial pressures of H_2_ were varied between 98 kPa H_2_ (at 0 kPa CO) and 49 kPa H_2_ (at 49 kPa CO) to maintain a constant availability of electron donors (98 kPa H_2_+CO) under all conditions (Fig. 1). Figure S6 shows time profiles of the electron balances under these conditions.

**Fig. 3.**
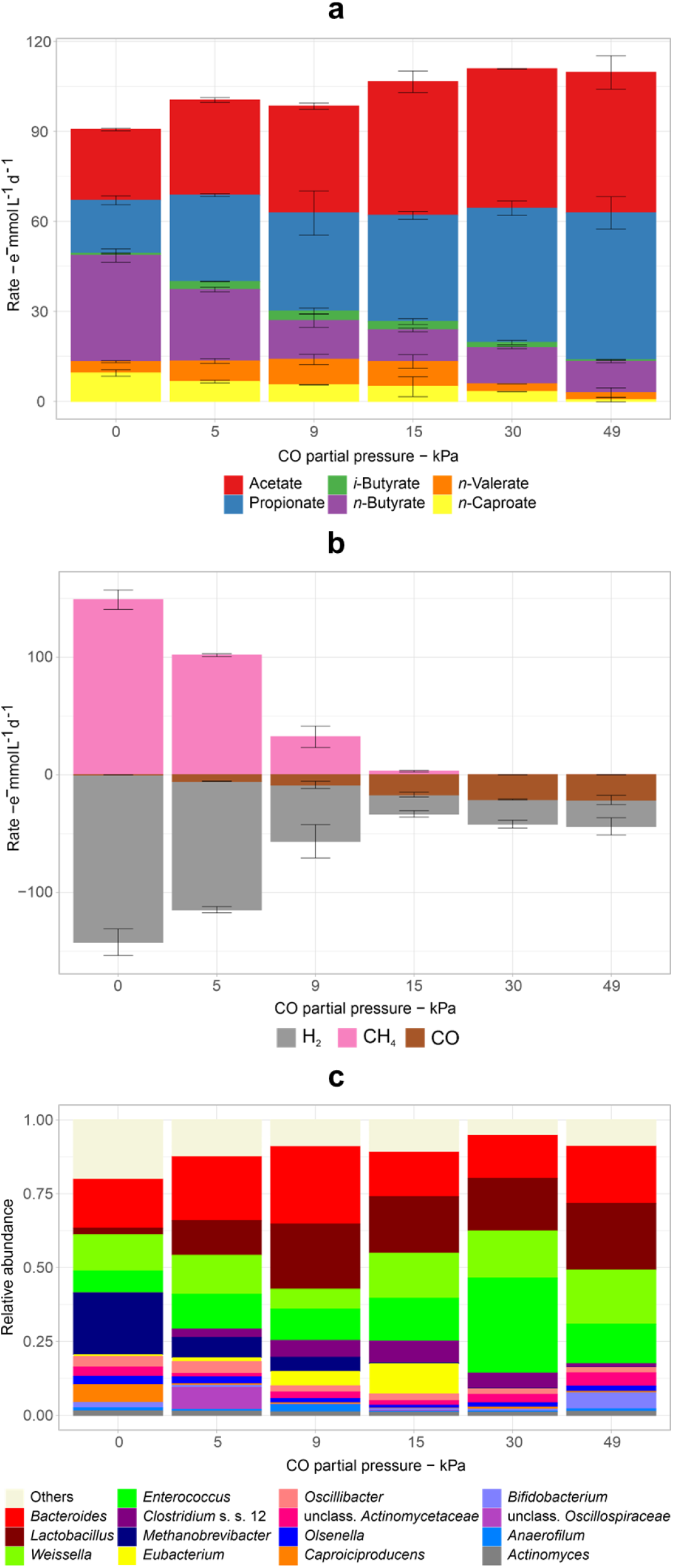
Effect of different partial pressures of CO on the carboxylate spectrum (**a**), gas production and consumption rates (**b**), and on the microbial community at the genus level (**c**). These tests were carried out with bottles inoculated with the adapted community. The 15 most abundant genera in the set are shown. Mean values of duplicate bottles are shown. Error bars indicate standard errors. Rates are in terms of electron equivalents. S.s.: sensu stricto.

As low as 5 kPa CO was enough to inhibit the electron flow to *n*-caproate and *n*-butyrate by 44% (Fig. 3a). On the other hand, production of acetate and propionate increased by 126% and 151%, respectively, when CO partial pressure increased from 0 to 49 kPa.

CO partial pressures above 9 kPa did not affect *n*-butyrate production but inhibited *n*-caproate formation, which dropped to almost zero at 49 kPa CO (Fig. 3a). *n*-Valerate production was only inhibited by more than 15 kPa CO. However, considering that *n*-valerate is produced from propionate [43], this apparent CO tolerance could just be the effect of increased propionate production.

Higher CO partial pressures favored the incorporation of H_2_ and CO into the carboxylate pool, although this effect became weaker with increasing CO pressure (Fig. 3a and b). At 5 and 9 kPa CO, the carboxylate pool increased likely due to partial inhibition of methanogenesis (Fig. 3b) and consequently higher H_2_/CO_2_ availability for acetogens. Nearly complete inhibition of methanogenesis by *Methanobrevibacter* was achieved at about 15 kPa CO, when CH_4_ accounted for less than 3% (3.1 ± 0.6 e^-^ mmol L^-1^ d^-1^) of the total electron sink (111 e^-^ mmol L^-1^ d^-1^) (Fig. 3b and c), similar to the study of Esquivel-Elizondo, et al. [44], in which 18 kPa CO completely inhibited methanogens in a mixed culture. An increase from 15 to 30 kPa CO improved H_2_ and CO consumption further from 16 ± 3 to 21 ± 3 e^-^ mmol H_2_ L^-1^ d^-1^ and from 17 ± 2 to 20.9 ± 0.4 e^-^ mmol CO L^-1^ d^-1^, whereas H_2_/CO consumptions were similar at 30 and 49 kPa CO (Fig. 3b).

Overall, increasing CO partial pressures (Fig. 3c) shaped the community consistently to what was observed previously when 49 kPa CO or 49 kPa H_2_ + 49 kPa CO were used (“Adapted community”, Fig. 2c). LAB, Actinobacteria (e.g., *Acidipropionibacterium* and *Actinomyces*), and *Bacteroides* were either unaffected or profited from increasing CO partial pressures, whereas *Caproiciproducens* was sensitive to CO (Fig. 3c). *Oscillibacter*, which generally had a comparably low abundance, was not inhibited by high CO partial pressures. This is relevant as *Oscillibacter* has previously been associated with *n*-valerate, *n*-caproate, and *n*-caprylate production as well as syngas consumption [45-47] and may be responsible for the formation of carboxylates with longer chains (C≥4) even at high CO pressures. A higher abundance of *Bifidobacterium* (a genus of lactate-producing actinobacteria) was observed at 49 kPa CO. Bacteria from this genus are selectively favored by high propionate concentrations [48], therefore, their higher relative abundance at high CO pressure could be a consequence of high propionate production rates (Fig. 3a) rather than of CO itself.

At high CO partial pressures, the rates of electrons routed to acetate (up to 47 ± 6 e^-^ mmol L^-1^ d^-1^ at 49 kPa CO) were close to the consumption rates of H_2_ and CO (44 e^-^ mmol L^-1^ d^-1^ H_2_+CO at 49 kPa CO) indicating predominant acetogenic activity. However, genera known to harbor acetogens (*Clostridium* sensu stricto 12 and *Eubacterium*) had higher relative abundances at intermediate CO partial pressures between 5 and 30 kPa (Fig. 3c).

### COMBINED METHANOGENESIS INHIBITION BY ETHYLENE AND CARBON MONOXIDE

Ethylene is a minor component of syngas [49] and an effective methanogenesis inhibitor at concentrations found in real syngas mixtures (in the order of 1%) [20]. Therefore, we studied its effect during co-fermentation of plant biomass when methanogenic activity was at its peak (at 0 and 5 kPa CO).

Overall, cultures with ethylene produced more carboxylates than cultures without ethylene (Fig. 4a). At 0 kPa CO (98 kPa H_2_) and at 5 kPa CO (93 kPa H_2_), ethylene addition increased the carboxylate production by 20% each. Ethylene showed no clear effect on the elongation of carboxylates and such increase was mainly due to the increased formation of acetate.

**Fig. 4.**
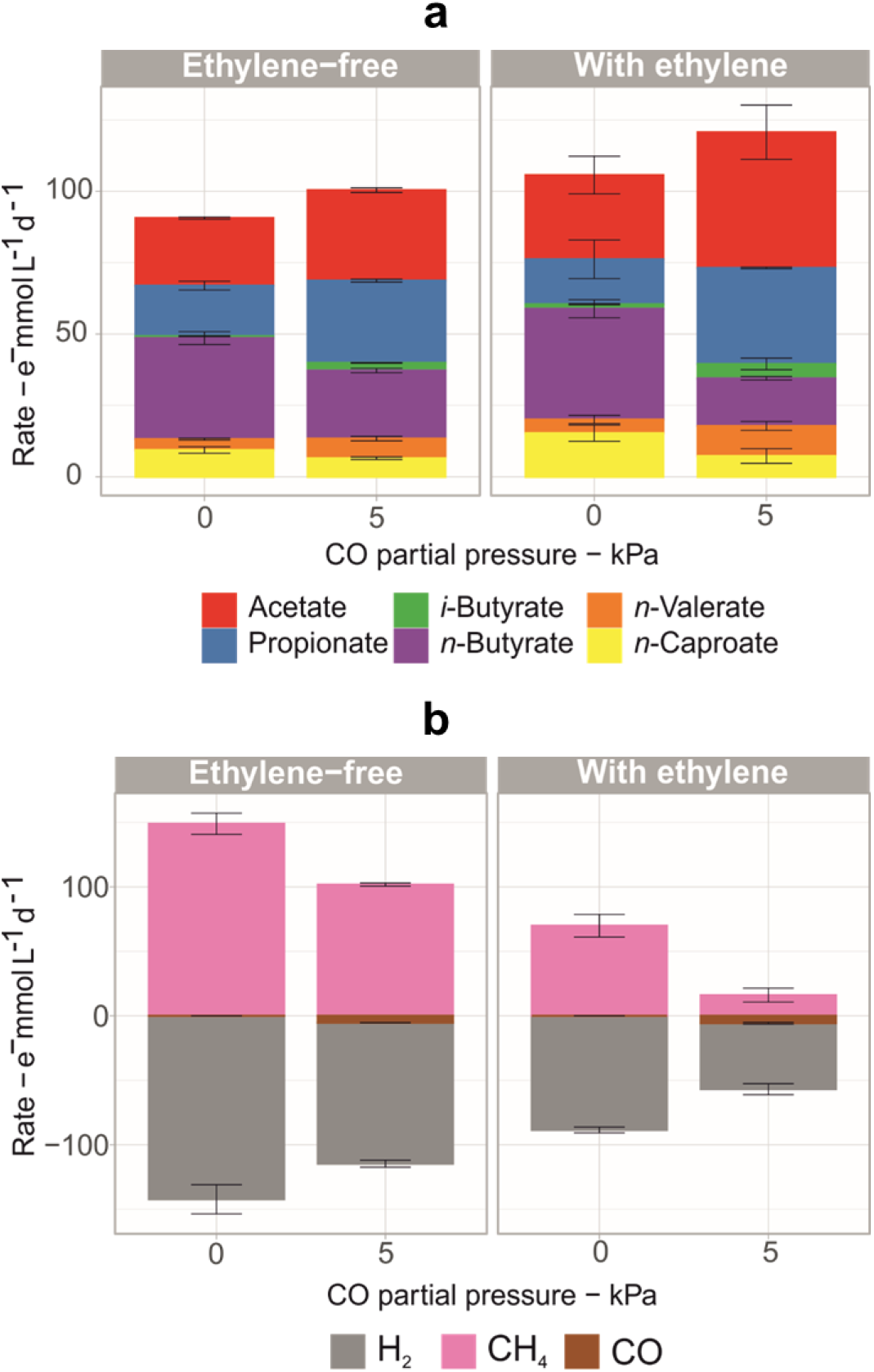
Effect of 1.5 kPa ethylene on the fermentation of corn silage with 0 kPa CO (98 kPa H_2_) and 5 kPa CO (93 kPa H_2_). Production or consumption rates of carboxylates (**a**) and gases (**b**) are shown in terms of electron equivalents. Error bars are standard errors. These tests were carried out with bottles inoculated with the adapted community.

Ethylene alone reduced methane production rates by about half (from 149 ± 8 to 69.9 ± 9 e^-^ mmol CH_4_ L^-1^ d^-1^), while 5 kPa CO alone reduced methane production rates by about one third (to 102 ± 1 e^-^ mmol CH_4_ L^-1^ d^-1^) (Fig. 4b). When CO and ethylene were applied together, CH_4_ production decreased 9-fold (to 16 ± 5 e^-^ mmol CH_4_ L^-1^ d^-1^). Ethylene had no effect on CO consumption but increased H_2_ consumption (excluding methanogenesis). Increase of H_2_ consumption due to ethylene presence was observed previously and was attributed to the increased H_2_ availability after inhibiting methanogens [20, 32]. No consumption of ethylene was observed.

Fig. 5 shows two benchmarks of the fermentation, i.e. carboxylate yields from the plant biomass and carbon fixation rates, at different CO partial pressures. The bottles with ethylene in combination with 5 kPa CO showed with 0.36 ± 0.07 g g_VS_^-1^ the highest carboxylate yield of this study (Fig. 5a). This yield is comparable to one of the highest yields achieved so far in continuous anaerobic fermenters fed with solid feedstock (without syngas and ethylene), namely 0.38 g g_VS_ ^-1^ [50]. The yield obtained with ethylene + 5 kPa CO was 44% higher than the yield achieved without syngas and 9% higher than the highest yield achieved with syngas containing 30 kPa CO.

**Fig. 5.**
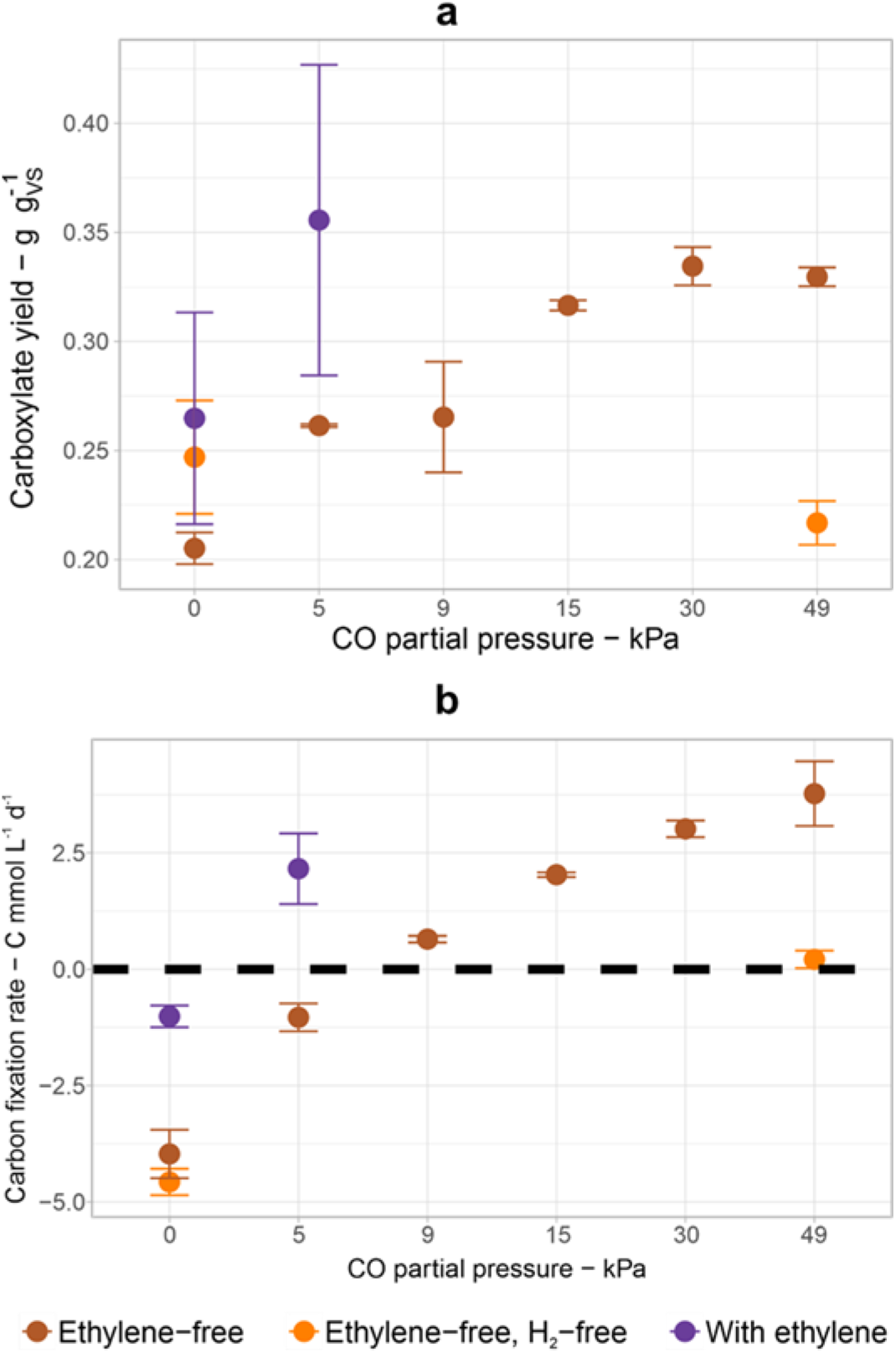
Carboxylate yields from biomass (**a**) and carbon fixation rates (**b**) of fermentations with different gas compositions with CO partial pressures between 0 and 49 kPa. Only results with the adapted community are shown.

The carbon fixation rates followed roughly the same trend as the carboxylate yields. Higher CO partial pressures allowed net carbon fixation up to a maximum of 3.0 ± 0.2 C mmol L^-1^ d^-1^ (equivalent to 132 ± 9 mg CO_2_ L^-1^ d^-1^) (Fig. 5b). This was achieved by inhibiting methanogenesis and by enhancing acetogenic activity, which in turn led to higher carboxylate production (Fig. 4). Cessation of carbon emissions was achieved when at least 9 kPa CO (without ethylene) or 5 kPa CO (with ethylene) was supplied. Fermentation with 5 kPa CO and ethylene was able to fix carbon at a rate of 2.2 ± 0.8 C mmol L^-1^ d^-1^, which was comparable to the carbon fixation rate of 2.03 ± 0.05 C mmol L^-1^ d^-1^ at 15 kPa CO (Fig. 5b).

When used separately, CO and ethylene are imperfect methanogenesis inhibitors. CO is not a selective inhibitor and although ethylene is [20], its inhibitory effect on archaeal hydrogenases can be bypassed by the expression of Fe-only hydrogenases [38]. When used together, small amounts of CO and ethylene had a synergistic effect in inhibiting hydrogenotrophic methanogens. Without methanogens that can misroute electrons from exogenous H_2_, anaerobic fermentation can be turned into a net carbon fixation process.

### THE ROLE OF FORMATE IN CARBON MONOXIDE TOLERANCE

Under most conditions, transient concentrations of up to 1 g L^-1^ formate were observed. In one pair of bottles with the autochthonous community in presence of syngas (49 kPa CO), formate accumulated in the first 10 days and then remained stable at about 2.14 ± 0.06 g L^-1^ (high formate, Figure S7). In this case, *n*-butyrate and *n*-caproate concentrations were clearly higher (1.7 ± 0.2 and 0.66 ± 0.09 g L^-1^, respectively) in comparison to a pair of bottles with low formate concentrations under the same conditions (0.5 ± 0.6 and 0.3 ± 0.1 g L^-1^, respectively) (low formate, Figure S7). From this observation, we suspected a relationship between formate and fermentative bacteria overcoming CO inhibition, similar to what has been observed in pure cultures of *A. woodii* [25]. We tested this hypothesis by adding 5 g L^-1^ formate at the beginning of the fermentation at 9 kPa CO. With 5 g L^-1^ formate, the production of *n*-butyrate and *n*-caproate at 9 kPa CO (3.3 ± 0.8 g L^-1^ and 0.9 ± 0.2 g L^-1^, respectively) was similar to that of the fermentations uninhibited by CO (3.1 ± 0.2 and 0.68 ± 0.08 g L^-1^, respectively) (Fig. 6).

**Fig. 6.**
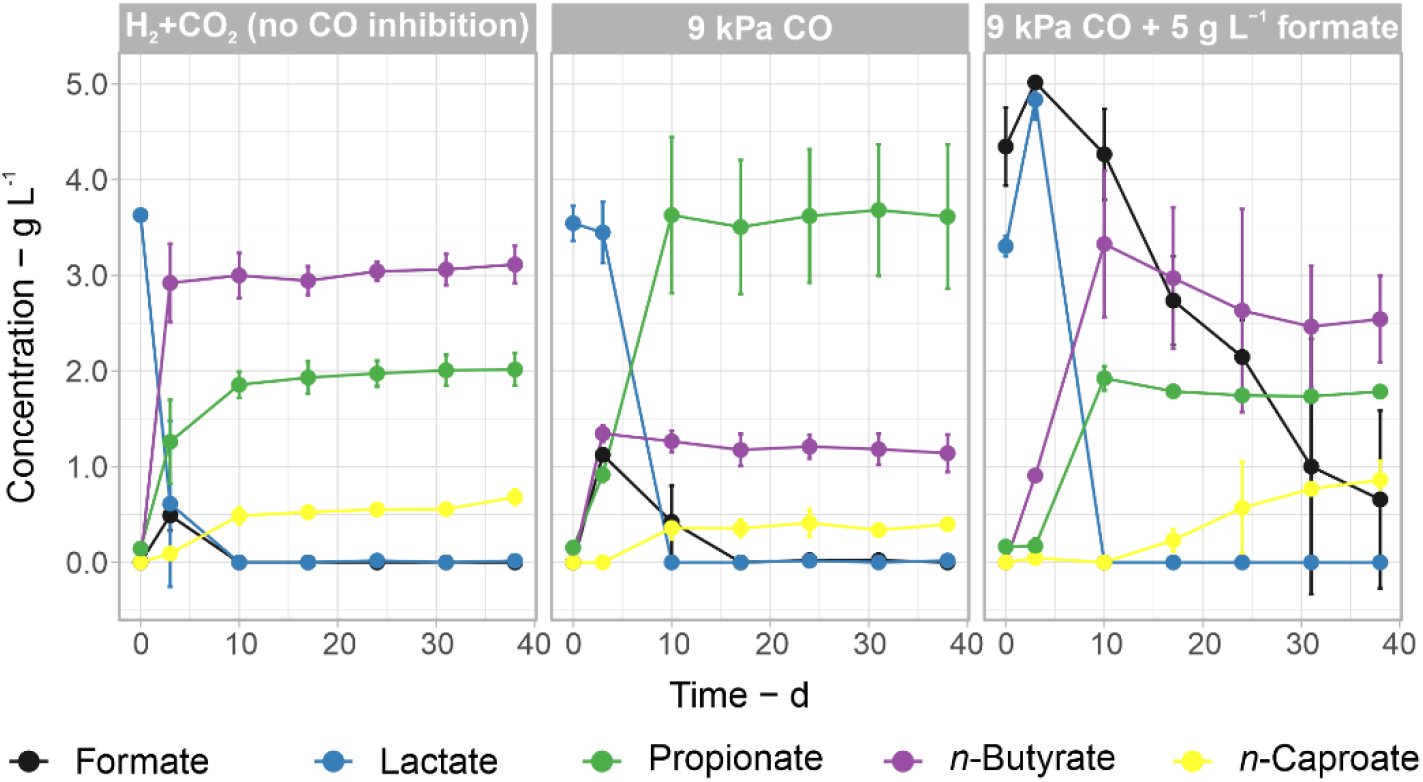
Fermentation profiles of the adapted community under a H_2_+CO_2_ headspace (uninhibited reference) and at 9 kPa CO with and without added formate. Mean values of duplicate bottles are shown, error bars are standard errors.

Based on the observations in Fig. 6 and Figure S7, we propose a theoretical model to explain how formate could help *n*-butyrate- and *n*-caproate-producing bacteria to overcome inhibition by CO (Fig. 7). These bacteria depend on hydrogenases to re-oxidize Fd_red_ (coupling it with H_2_ formation) or to regenerate their NAD(P)H pools. NAD(P)H is required for the elongation cycles with the acyl-CoA and 3-hydroxy-acyl-CoA dehydrogenases (ACAD and 3-HACAD, respectively). On the other hand, formate dehydrogenases (Fdh) can be found in many acidogenic bacteria and these enzymes are assumed to be less sensitive to CO than hydrogenases [51]. Thus, *n*-butyrate and *n*-caproate producers that have an Fdh can couple formate oxidation with NAD^+^ reduction and bridge the gap left by hydrogenases inhibited by CO.

**Fig. 7.**
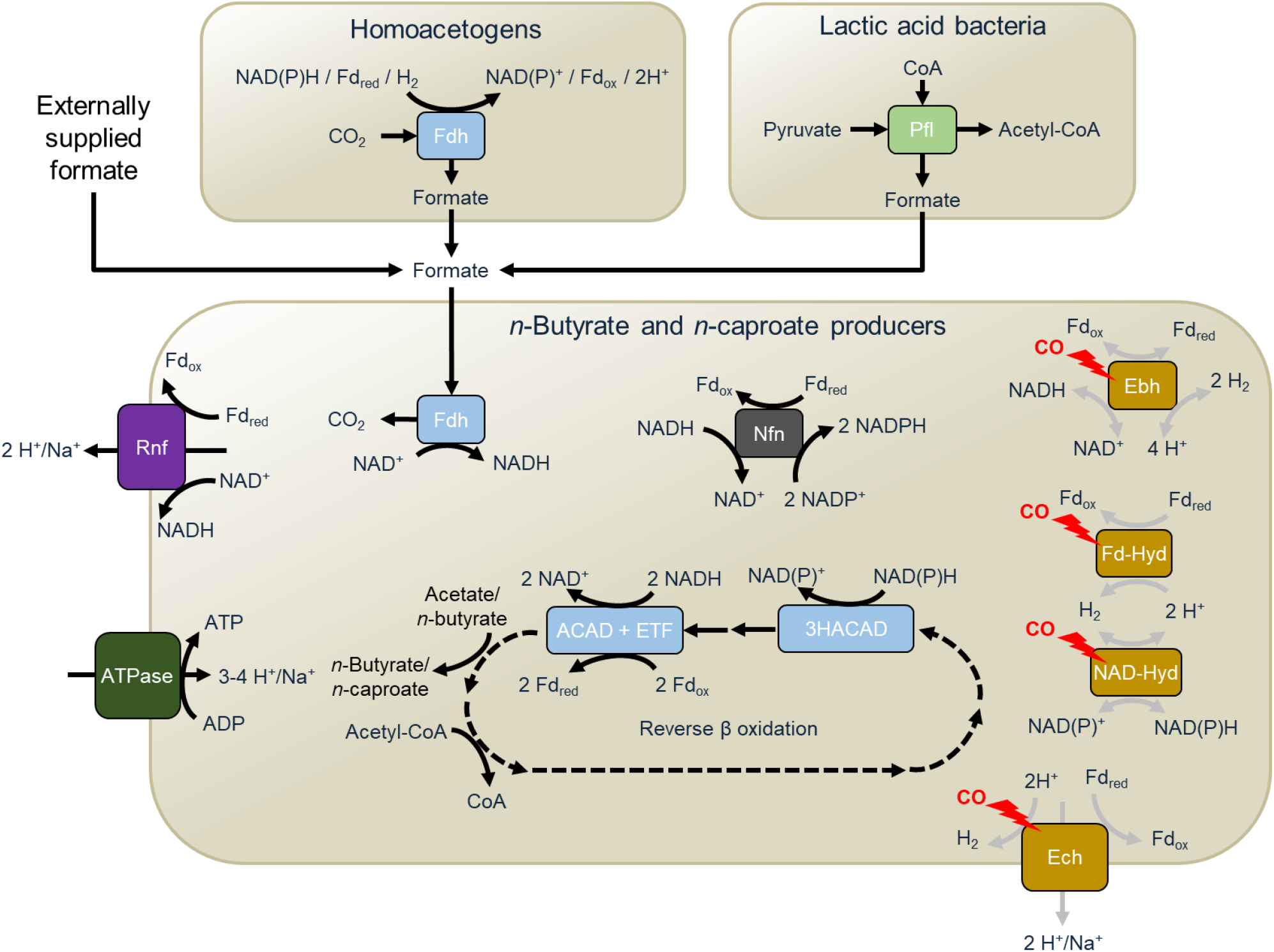
Theoretical model of formate-induced CO tolerance in *n*-butyrate and *n*-caproate producers performing reverse β-oxidation. The CO-inhibited enzymes in the lower box are typically involved in energy conservation in anaerobic acidogenic bacteria that lack cytochromes. Chain-elongating bacteria can compensate for this inhibition by oxidizing formate, thereby providing reduction equivalents needed for reverse β-oxidation. Dashed arrows represent sequential metabolic steps and grey arrows represent CO-inhibited processes. Abbreviations: 3-HACAD, 3-hydroxy-acyl-CoA dehydrogenase; ACAD, acyl-CoA dehydrogenase; Ebh, electron bifurcation hydrogenase; Ech, energy-conserving hydrogenase; ETF, electron-transferring flavoproteins A and B; Fdh, formate dehydrogenase; Fd-Hyd, Fd-dependent hydrogenase; NAD-Hyd, NAD-dependent hydrogenase; Nfn, NAD(P)^+^ transhydrogenase; Pfl, pyruvate formate lyase.

Formate is an extracellular electron carrier between anaerobic microorganisms such as methanogens, acetogens, and sulfate-reducing bacteria [52, 53]. Under the conditions used here, we expect endogenous formate production to occur mainly in two different ways (Fig. 7): (i) during the conversion of pyruvate to acetyl-CoA via pyruvate formate lyase (Pfl) often found in LAB [41] and (ii) as an intermediate in the metabolism of acetogens after the fixation of one CO_2_ molecule with an electron pair (in the form of H_2_, NADH, or Fd^-2^) via Fdh in the methyl branch of the Wood-Ljungdahl pathway [54, 55]. According to the UniProt database [56], Fdh is predicted from the genomes of *C. butyricum* (*Clostridium* sensu stricto 1), *Megasphaera elsdenii, Eubacterium limosum*, and *C. luticellarii* (*Clostridium* sensu stricto 12). However, some other well-known acidogens such as *C. kluyveri* and *C. tyrobutyricum* (both belonging to *Clostridium* sensu stricto 12) do not have annotated genes for Fdh.

Figure S8 presents the community compositions at the genus level for the cultures with inhibition or tolerance to CO. Higher relative abundances of *Clostridium* sensu stricto 1 were observed in the fermentation with the autochthonous community under the conditions CO+formate and H_2_+CO_2_ (no CO inhibition). In the adapted community, the presence of formate caused the absence of *Bacteroides*, which in contrast accounted for about 25% of the bacteria in the bottles without formate, and increased the share of less abundant genera (grouped in “others”), in similarity to uninhibited bottles. 16S rRNA amplicon sequencing does not provide direct information on the presence or absence of the Fdh gene in bacteria. Thus, we were not able to investigate links between Fdh-containing bacteria and formate-induced CO tolerance. Therefore, the proposed metabolic mechanism in Fig. 7 should be further tested experimentally. Preferably, this should be done in less complex systems such as co-cultures or single cultures, while monitoring the expression of the respective genes.

Besides, there are other conceivable explanations for how formate restores the production of *n*-butyrate and *n*-caproate in CO-inhibited cultures. It is possible that formate favors *n*-butyrate and *n*-caproate producers by selectively inhibiting propionate producers that compete for lactate and sugars. Direct conversion of formate to *n*-butyrate could be another explanation. However, this conversion is energetically unfavorable [57]. To the best of our knowledge, pure cultures of acetogenic bacteria produce at best trace amounts of *n*-butyrate (and *n*-caproate) from formate. For instance, *E. limosum* can grow on formate but without *n*-butyrate production [58].

## CONCLUSION

CO toxicity and inhibition of methanogens were the most important factors influencing the carboxylate production. Increasing CO partial pressures completely reshaped the microbial community and shifted the product spectrum from C≥4 carboxylates and methane to acetate and propionate. H_2_ in the headspace had limited effects. When H_2_ was present, it favored the growth of hydrogenotrophic methanogens, inhibited some bacterial genera commonly associated with lignocellulose degradation, and slowed biomass decomposition. From a sustainability perspective, a syngas composition with low partial pressures of CO and ethylene and high partial pressure of H_2_ was particularly interesting since it showed a synergistic effect in inhibiting hydrogenotrophic methanogens and achieving net carbon fixation via acetogenesis. This syngas mixture yielded 44% more carboxylates than a conventional fermentation (with a N_2_/CO_2_ headspace) despite slower biomass degradation rates, hence reducing the dependence of anaerobic fermentation on biomass availability. From a process engineering perspective, costs could be saved by not having to remove ethylene from real syngas mixtures. Nevertheless, to achieve high yields of longer-chain carboxylates (C≥4), we recommend testing the concept by operating the fermentation as a continuous process. In this way, chain-elongating bacteria that cope well with both CO and complex feedstocks could be selected.

Under most conditions, as little as 5 kPa CO was sufficient to hinder *n*-caproate and *n*-butyrate production. However, this inhibition was not observed in cultures with CO in which formate concentration remained above 2 g L^-1^. To the best of our knowledge, formate-induced tolerance to CO has not yet been reported, neither for mixed cultures nor for *n*-butyrate- and *n*-caproate-producing pure cultures. We postulate that formate could help fermentative bacteria maintain their NAD(P)H pool via formate dehydrogenase, thus bridging the gap left by hydrogenases inhibited by CO. Further experiments are needed to test this hypothesis. If true, this feature could be exploited in designing bioelectrochemical systems with CO or in fermentation technologies based on C1 substrates.

## Supplementary Information

Additional file 1. Mineral medium preparation, Further details on the experimental setup, and Inoculation with a syngas-adapted community. Table S1. Mineral medium composition. Table S2. Conversion factors. Figure S1. Concentration of organic acids and electron balances in the abiotic controls. Figure S2. Composition of the syngas-adapted community used as inoculum. Figure S3. Concentration of organic acids and electron balances during the fermentation of corn silage + syngas (H_2_/CO/CO_2_ ratio of 49:49:24 kPa) depending on the inoculation with the syngas-adapted community. Figure S4. Carbon fixation rates achieved with the autochthonous corn silage community and with the inoculated (adapted) community at different components of syngas. Figure S5. Solids degradation relative to the abiotic controls (in percentage points, p.p.) during the fermentation of corn silage in the presence of different gases and with different inocula. Figure S6. Electron balance kinetics for fermentations of corn silage in the presence of different partial pressures of CO. Figure S7. Fermentation profiles of the autochthonous community under a H_2_+CO_2_ headspace (uninhibited reference) and under a syngas headspace (49 kPa CO) with low and high formate accumulation. Figure S8. Composition of the autochthonous community and the syngas-adapted community depending on CO inhibition and formate availability.

## DECLARATIONS

### Ethics approval and consent to participate

Not applicable.

### Consent for publication

Not applicable.

### Availability of data

The raw sequence reads generated during the current study are available in the European Nucleotide Archive (ENA) under the accession PRJEB49567 (http://www.ebi.ac.uk/ena/data/view/PRJEB49567).

### Authors’ contributions

FCFB, SK, AN, HS, and LV conceptualized the study and reviewed the manuscript. FCFB and LV developed the methodology, performed the experiments, and analyzed the data. FCFB prepared the original draft. SK, AN, and HS supervised the project and supported the data analysis. All authors have given approval to the final version of the manuscript.

## Acknowledgments

We thank Ute Lohse for technical assistance in library preparation for MiSeq amplicon sequencing and Pascal Gorenflo for his support in the HPLC operation and maintenance.

