## Additional file 1 for "Formate-induced CO tolerance and innovative methanogenesis inhibition in co-fermentation of syngas and plant biomass for carboxylate production"

MINERAL MEDIUM PREPARATION. All medium ingredients (Table S1), except vitamins and cysteine, were mixed under oxic conditions. After adjusting the pH to 6.0 with 4 M NaOH, the mixture was stirred (700 rpm) in an anaerobic chamber (1-5% $H_2$ with the rest as $N_2$) for at least 3 hours to be make it anoxic. The anoxic solution was transferred to a glass bottle and capped and autoclaved at 121°C for 20 minutes. Stock solutions with vitamins and with cysteine were prepared separately and were only mixed with the rest of the ingredients on the day of the start of the batch experiment inside the anaerobic chamber. The vitamin and cysteine stock solutions were stored in serum bottles sealed with butyl rubber stoppers and aluminum crimps. The concentrated 30 g $L^{-1}$ cysteine hydrochloride solution was prepared with anoxic, deionized water inside an anaerobic chamber and was made sterile by autoclaving at 121°C for 20 minutes. The 1000-fold concentrated vitamin stock solution was sterilized with a nylon 0.22 μm syringe filter and then made anoxic with at least 15 pressurization-depressurization cycles with pure $N_2$ lasting 2 min each.

FURTHER DETAILS ON THE EXPERIMENTAL SETUP. The experiments described in Fig. 1 were divided into three batches lasting between 31 and 38 d depending on the carboxylate production. The first batch lasted 31 d and compared different starting communities (the autochthonous and the syngas-adapted communities) as well as the way of inoculating the adapted community (with washed cells, 10 vol % reactor broth, or 100 vol % reactor broth). This batch was realized under a syngas atmosphere with 49 kPa $H_2$, 49 kPa CO, and 24 kPa $CO_2$. The second batch lasted 35 d and compared the effects of the presence or absence of $H_2$ and CO on the autochthonous community and on the adapted community. The third batch lasted 38 d and tested the adapted community under intermediate partial pressures of CO (from 0 to 30 kPa). During this batch, the effect of 1.5 kPa ethylene (at 0 and 5 kPa CO) and the effect of the addition of 5 g $L^{-1}$ formate (at 9 kPa CO) were also tested.

**Table S1.** Mineral medium composition.

| | |
|---|---|
| Concentration of major components - g $L^{-1}$ | |
| $NH_4Cl$ | 1.61 |
| $KH_2PO_4$ | 13.6 |
| NaCl | 2.0 |
| NaOH (approx.) | >1.0 (correction to pH 6.0) |
| Concentration of minor components - mg $L^{-1}$ | |
| $MgCl_2 \times 6\ H_2O$ | 54 |
| $CaCl_2 \times 2\ H_2O$ | 65 |
| Resazurin | 0.5 |
| $FeCl_2 \times 4\ H_2O$ | 1.5 |
| Cysteine hydrochloride | 30 |
| Concentration of trace elements - µg $L^{-1}$ | |
| $CuCl_2 \times 2\ H_2O$ | 2.0 |
| $CoCl_2 \times 6\ H_2O$ | 190 |
| $MnCl_2$ | 100 |
| $Na_2MoO_4 \times 2\ H_2O$ | 36 |
| $NiCl_2 \times 6\ H_2O$ | 24 |
| $Na_2WO_4 \times 2\ H_2O$ | 20 |
| $Na_2SeO_3 \times 5\ H_2O$ | 3.0 |
| $ZnCl_2$ | 70 |
| $H_3BO_3$ | 6.0 |
| Concentration of vitamins - µg $L^{-1}$ | |
| Biotin | 20 |
| Folic acid | 20 |
| Pyridoxine | 100 |
| Thiamine | 50 |
| Riboflavin | 50 |
| Niacin | 50 |
| Calcium pantothenate | 50 |
| Cobalamin | 20 |
| p-Aminobenzoic acid | 80 |
| Lipoic acid | 50 |

**Table S2.** Conversion factors.

| Compound (formula) | Molar mass (g $mol^{-1}$) | mol C $mol^{-1}$ | mol $e^-$ $mol^{-1}$ |
|---|---|---|---|
| Biomass ($C_1H_{1.8}O_{0.5}N_{0.2}$) | 24.6 | 1.0 | 4.2 |
| Formic acid/formate ($C_1H_2O_2$) | 46.0 | 1.0 | 2.0 |
| Acetic acid/acetate ($C_2H_4O_2$) | 60.0 | 2.0 | 8.0 |
| Ethanol ($C_2H_6O$) | 46.0 | 2.0 | 12.0 |
| Propionic acid/propionate ($C_3H_6O_2$) | 74.0 | 3.0 | 14.0 |
| Lactic acid/lactate ($C_3H_6O_3$) | 90.0 | 3.0 | 12.0 |
| *n*-Butyric acid/*n*-butyrate ($C_4H_8O_2$) | 88.0 | 4.0 | 20.0 |
| *i*-Butyric acid/*i*-butyrate ($C_4H_8O_2$) | 88.0 | 4.0 | 20.0 |
| *n*-Valeric acid/ *n*-valerate ($C_5H_{10}O_2$) | 102.1 | 5.0 | 26.0 |
| *n*-Caproic acid/ *n*-caproate ($C_6H_{12}O_2$) | 116.1 | 6.0 | 32.0 |
| $H_2$ | 2.0 | 0.0 | 2.0 |
| $CO_2$ | 44.0 | 1.0 | 0.0 |
| CO | 28.01 | 1.0 | 2.0 |
| $CH_4$ | 16.0 | 1.0 | 8.0 |

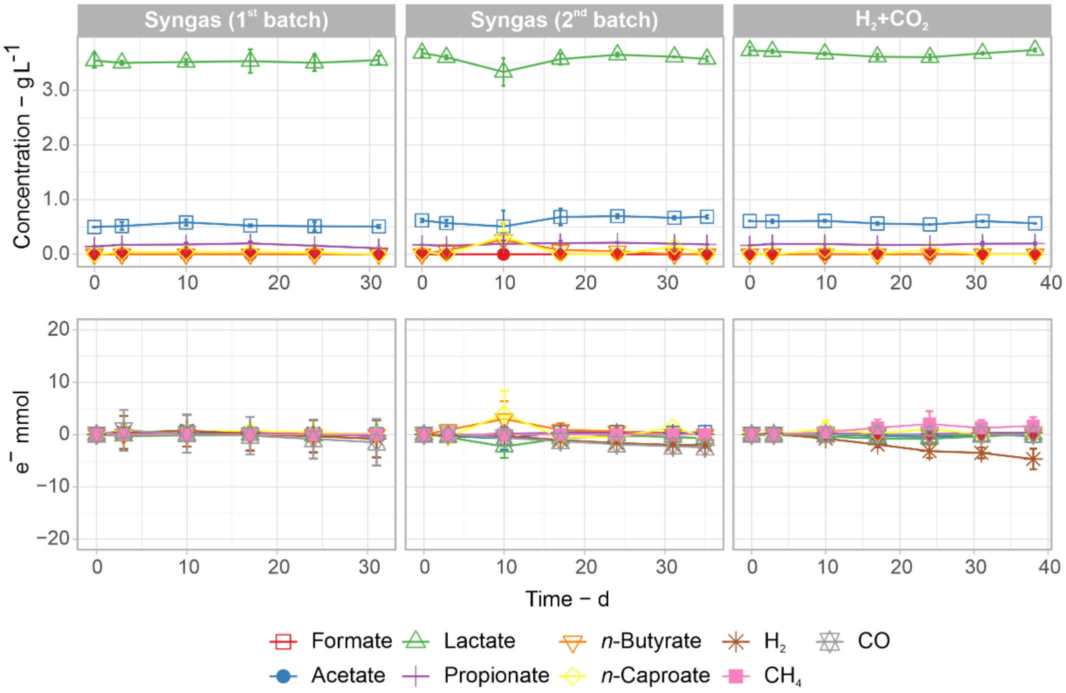

**Figure S1.** Concentration of organic acids and electron balances in the abiotic controls. Mean values of duplicate bottles are shown. Error bars indicate standard errors.

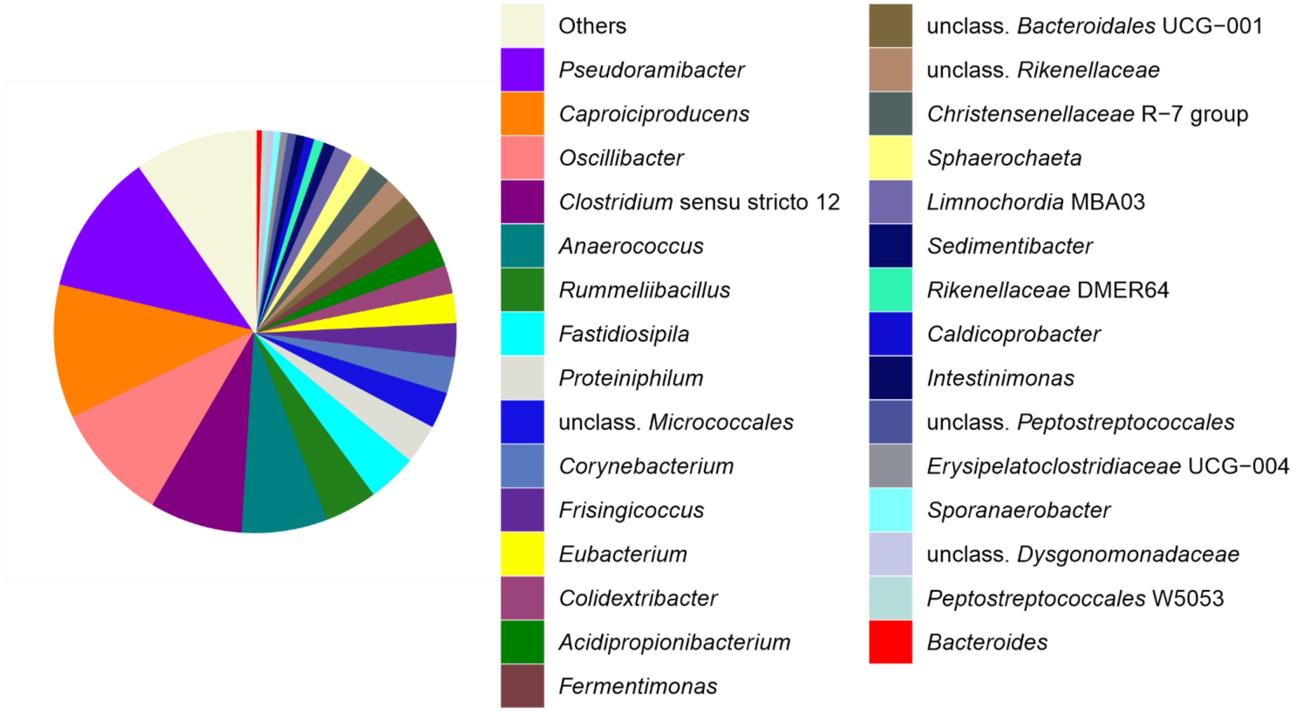

**Figure S2.** Composition of the syngas-adapted community used as inoculum. The 30 most abundant genera are shown.

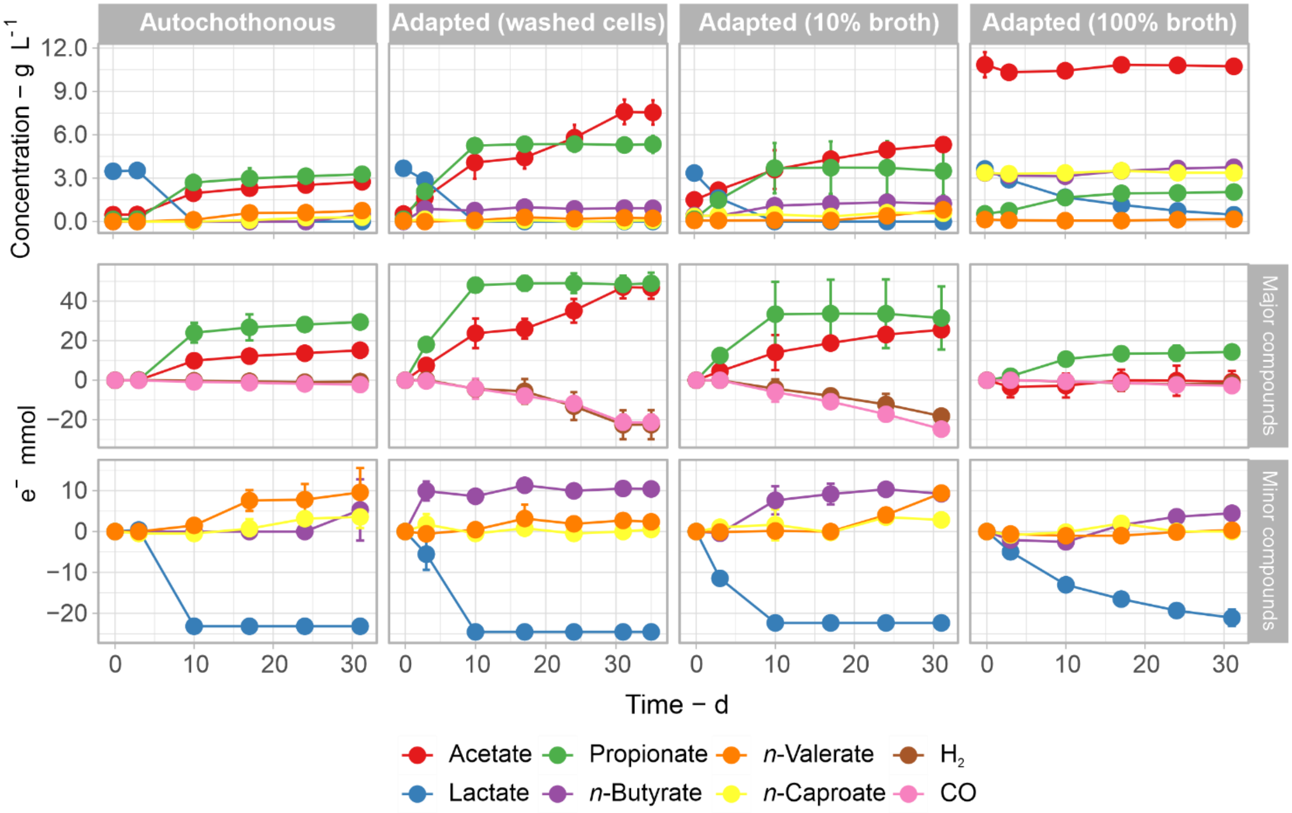

**Figure S3.** Concentration of organic acids and electron balances during the fermentation of corn silage + syngas ($H_2/CO/CO_2$ ratio of 49:49:24 kPa) depending on the inoculation with the syngas-adapted community (see Figure S2). From left to right: autochthonous community (corn silage community only), inoculation with the adapted community (washed cells), inoculation with 10 vol% reactor broth, and inoculation with 100 vol% reactor broth. Mean values of duplicate bottles are shown. Error bars indicate standard errors.

INOCULATION WITH A SYNGAS-ADAPTED COMMUNITY. Regardless of the initial microbial community, production of carboxylates with more than three carbon atoms (i.e. *n*-butyrate, *n*-valerate, and *n*-caproate) remained less than 1.5 g $L^{-1}$ (Figure S3). Besides, no methane formation was observed.

Cultures that contained only the autochthonous community of corn silage presented a longer lag phase than bottles inoculated with the adapted community, as evidenced by the late start of lactate consumption. Still, the autochthonous community started to slowly consume CO in the middle of the batch.

In comparison to the fermentations with the corn silage community alone, inoculation with the adapted community via washed cells or via the addition of 10 vol% reactor broth caused higher production of acetate and propionate. In addition, the highest $H_2$ and CO consumption was also observed in these two sets.

Originally, the reactor broth used for the inoculation contained a high concentration of carboxylates (ca. 11 g $L^{-1}$ acetate, 4.5 g $L^{-1}$ C4 carboxylates, and 2.6 g $L^{-1}$ *n*-caproate). Therefore, when 100 vol% of the enrichment reactor broth was used as inoculum, the carboxylates present in the broth strongly inhibited the fermentation of corn silage and syngas, and only a slow conversion of the lactate from corn silage into propionate was observed. The carboxylates from the broth that entered the bottles via inoculation were neither consumed nor converted into alcohols.

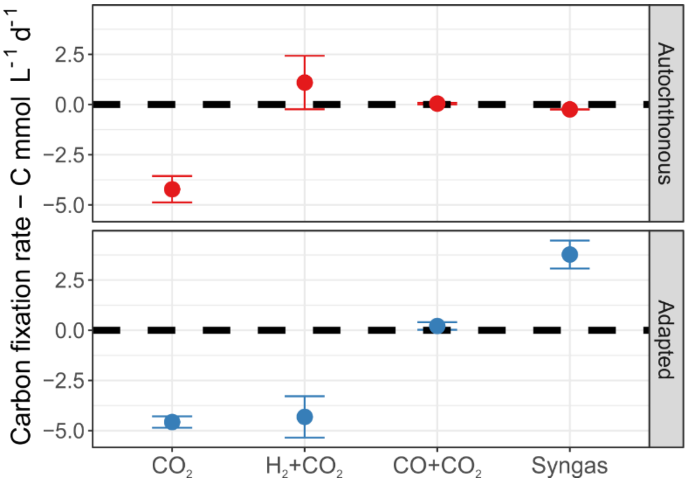

**Figure S4.** Carbon fixation rates achieved with the autochthonous corn silage community and with the inoculated (adapted) community at different components of syngas. Mean values of duplicate bottles are shown. Error bars indicate standard errors.

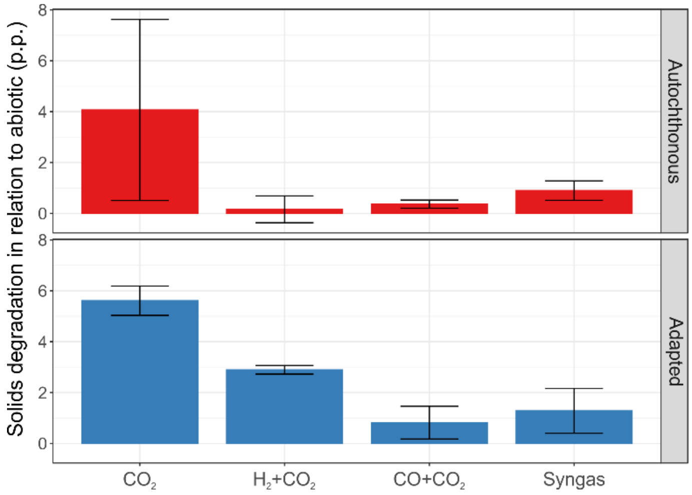

**Figure S5.** Solids degradation relative to the abiotic controls (in percentage points, p.p.) during the fermentation of corn silage in the presence of different gases and with different inocula. Mean values of duplicate bottles are shown. Error bars indicate standard errors.

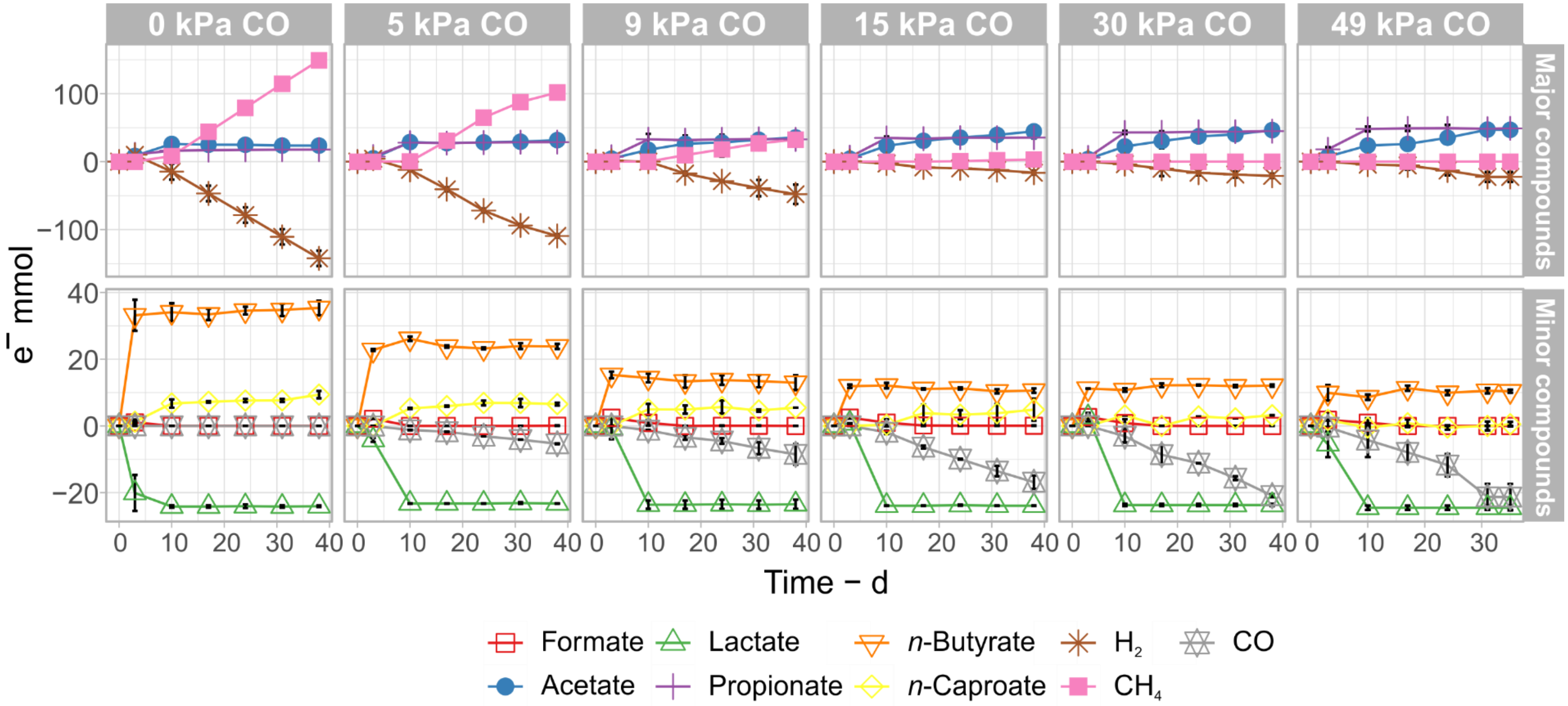

**Figure S6.** Electron balance kinetics for fermentations of corn silage in the presence of different partial pressures of CO. Cultures inoculated with the adapted community were used for this experiment. Mean values of duplicate bottles are shown. Error bars indicate standard errors.

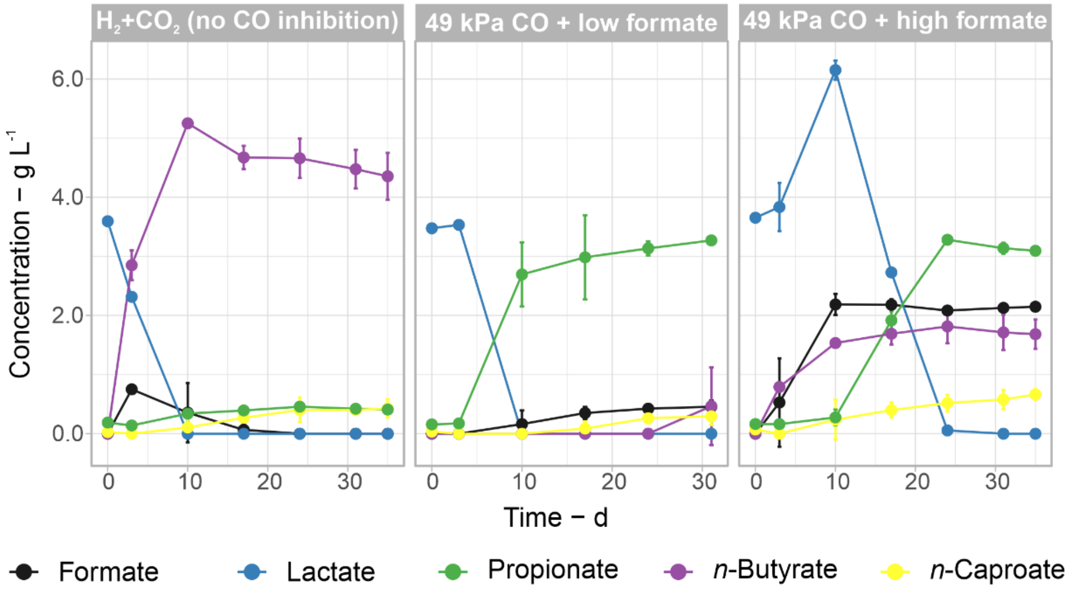

**Figure S7.** Fermentation profiles of the autochthonous community under a $H_2$+$CO_2$ headspace (uninhibited reference) and under a syngas headspace (49 kPa CO) with low and high formate accumulation. Mean values of duplicate bottles are shown. Error bars indicate standard errors.

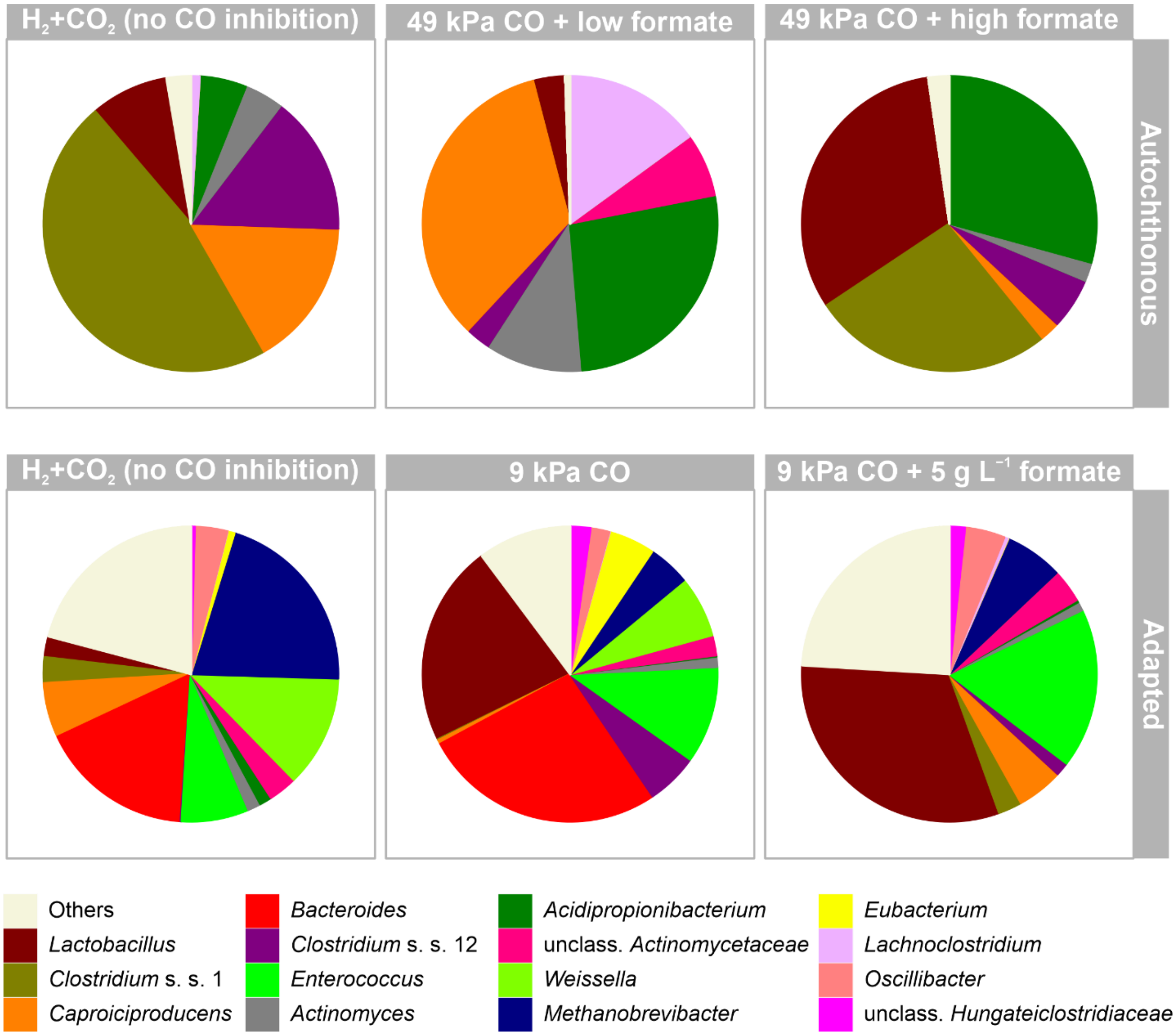

**Figure S8.** Composition of the autochthonous community and the syngas-adapted community depending on CO inhibition and formate availability. The 15 most abundant genera in the set are shown. S.s.: sensu stricto.
